# Highly functional Cellular Immunity in SARS-CoV-2 Non-Seroconvertors is associated with immune protection

**DOI:** 10.1101/2021.05.04.438781

**Authors:** Athina Kilpeläinen, Esther Jimenez-Moyano, Oscar Blanch-Lombarte, Dan Ouchi, Ruth Peña, Bibiana Quirant-Sanchez, Anna Chamorro, Ignacio Blanco, Eva Martínez-Caceres, Roger Paredes, Lourdes Mateu, Jorge Carrillo, Julià Blanco, Christian Brander, Marta Massanella, Bonaventura Clotet, Julia G. Prado

## Abstract

The role of T cells in the control of SARS-CoV-2 infection has been underestimated in favor of neutralizing antibodies. However, cellular immunity is essential for long-term viral control and protection from disease severity. To understand T-cell immunity in the absence of antibody generation we focused on a group of SARS-CoV-2 Non-Seroconvertors (NSC) recovered from infection. We performed an immune comparative analysis of SARS-CoV-2 infected individuals stratified by the absence or presence of seroconversion and disease severity. We report high levels of total naïve and low effector CD8+ T cells in NSC. Moreover, polyfunctional Nucleocapsid (NP)-specific CD8+ T-cell responses, as well as reduced levels of T-cell activation monitored by PD-1 and activation-induced markers, were distinctive immunological traits in NSC. Longitudinal data support the stability of the NSC phenotype over three months. Our results implicate highly functional SARS-CoV-2 Spike and NP T-cell responses with low immune activation in protection from disease severity in the absence of seroconversion.

**SUMMARY:** To understand SARS-CoV-2 specific T-cell immunity in the absence of seroconversion, we characterized immunological features of Non-Seroconvertors recovered from infection. Highly functional specific T-cell responses and low immune activation were determinants of immune protection from severe disease.

## INTRODUCTION

The COVID-19 pandemic is caused by SARS-CoV-2, the newest coronavirus crossing into the human population. The pandemic accounts for millions of infected people and more than 2 million deaths worldwide. Despite the scientific success in rapidly generating SARS-CoV-2 vaccines, the number of infected people and the burden on healthcare systems continues (Chang et al., 2020). Consequently, continuous characterization of the functional features of immune protection for control and prevention of SARS-CoV-2 infection are urgently needed.

Disease outcome of SARS-CoV-2 infection is associated with a wide degree of interindividual heterogeneity. These divergences may be associated with the level of immunocompetence and the tight balance between immune control and immunopathogenesis after infection (Blanco-Melo et al., 2020). Indeed, beneficial and detrimental aspects of the immune responses elicited by SARS-CoV-2 infection have been described. As detrimental aspects, secondary multi-organ complications and persistent symptomatology for months in approximately 10% of the total infected people or “long COVID” (Marshall, 2020) have been associated with the persistence of inflammatory profiles and tissue damage. As beneficial aspects, cellular and humoral immune responses have been linked to protection from disease severity against other coronaviruses such as SARS-CoV-1 (Li et al., 2006) and the induction of B and T cell memory responses has been described in most of the individuals recovered from SARS-CoV-2 infection (Dan et al., 2021).

Disease outcomes have been correlated with early signatures of soluble meditators including growth factors, type-2/3 cytokines, type-1/2/3 cytokines, and chemokines in SARS-CoV-2 infection (Lucas et al., 2020). Cellular perturbations and immune profiles have been related to disease severity (Kuri-Cervantes et al., 2020; Wilk et al., 2020; Mathew et al., 2020). Also, neutralizing antibodies have been shown to predict severity and survival (Garcia-Beltran et al., 2021). However, high neutralizing antibody titers do not positively correlate with less severe clinical outcome (Trinité et al., 2021; Rydyznski Moderbacher et al., 2020).

Antiviral CD4+ and CD8+ T-cell responses are key in the natural control of viral infections (Noel et al., 2016; Pereyra et al., 2010; Kiepiela et al., 2007). Emerging data from SARS-CoV-2 animal models support the role of T-cell immunity as a correlate of protection from infection (McMahan et al., 2021) and virus-specific CD4+ and CD8+ T-cell responses are considered key players in the resolution and long-term protection from infection (Peng et al., 2020). The presence of SARS-CoV-2 CD4+ and CD8+ T-cell responses up to 6 months after infection in mild to moderate clinical course and persistent immunological alterations in the memory compartment have been described (Breton et al., 2021; Dan et al., 2021). Moreover, protection through pre-existent SARS-CoV-2 cross-reactive CD4+ T cell responses to other coronaviruses in a fraction of individuals has been proposed as a mechanism limiting disease severity (Mateus et al., 2020; Sekine et al., 2020). These and other studies support the relevance of cellular immunity in the control and prevention of SARS-CoV-2 infection.

Phenotypic and functional characterization of cellular immunity against SARS-CoV-2 in the absence of immune pathogenesis remains limited but is essential to identify the determinants of immune protection from infection and disease severity. To gain deeper insights of cellular immune protection, we focused on a group of SARS-CoV-2 Non-Seroconvertors (NSC), defined as individuals with confirmed SARS-CoV-2 infection in the absence of antibody generation and compared their total and virus-specific T cells to individuals who seroconverted. Lack of seroconversion in SARS-CoV-2 infection has been demonstrated in 2-17% of convalescent individuals (Schulien et al., 2021; Schwarzkopf et al., 2021; Sekine et al., 2020; Staines et al., 2021) and cellular responses have been observed in 41 to 78% of these individuals (Schwarzkopf et al., 2021; Sekine et al., 2020). Currently, the functional characterization of T cell responses in NSC remains limited due to the low numbers of individuals identified, lack of confirmation of SARS-CoV-2 infection by PCR, or specific focus in CD8+ T cells. To fill this gap, we characterized SARS-CoV-2-specific CD4+ and CD8+ T cell responses in NSC in terms of the cellular landscape, function, and activation profile and compared these to responses detected in recovered individuals that seroconverted stratified by the degree of viral neutralizing activity and SARS-CoV-2 disease severity. Our analyses identify differential immunological traits in NSC including skewed CD8+ T cell distribution towards an increase in naïve populations and polyfunctional Nucleocapsid protein (NP) SARS-CoV-2-specific CD8+ T cell responses in the absence of overt immune activation. Our data highlight a non-redundant and essential contribution of T-cell immunity against SARS-CoV-2 in immune protection from severity in the absence of humoral immunity. Our current analyses of SARS-CoV-2 T-cell immunity provides information required for clinical monitoring and guiding future immune interventions.

## MATERIALS AND METHODS

### Study participants

The KING extension cohort is composed of 336 SARS-CoV-2 infected individuals, including 120 individuals who required hospitalization, 22% of whom with severe or critical SARS-CoV-2 infection (**Fig 1A**). For the present study we identified participants (N=52) (median age 47 years, 62% female, 38% male) across Non-Seroconvertors (NSC, n=16), Low-neutralizers (LN, N=16) and Convalescent (Conv, N=15) (**Fig 1A**). Also, we obtained samples from healthy donors from the Catalan Blood and Tissue Bank before 2019 as controls (Donor, N=5). The NSC represent 4.8% of the King extension cohort and were defined as PCR-confirmed SARS-CoV-2 infected individuals with no detectable SARS-CoV-2 IgG, IgA and IgM by two independent ELISAs (in-house and IMD). Moreover, the lack of seroconversion was confirmed in 10 NSC a median of 94.5 days after symptom onset and longitudinal follow-up was available for 6 of them. The LN were defined as having detectable anti-SARS-CoV-2 antibodies with neutralizing titers <500 IC50 values measured by pseudovirus neutralization assay. The study groups were stratified by the presence or absence of seroconversion, viral neutralizing titers and SARS-CoV-2 disease severity. The clinical characteristics of participants are summarized in **Table 1**. All methods and experimental protocols of the study were approved by the Ethics Committee of Hospital Germans Trias i Pujol (PI-20-217). All the study subjects provided their written informed consent to participate. The study was conducted according to the principles expressed in the Declaration of Helsinki.

**Figure 1.**
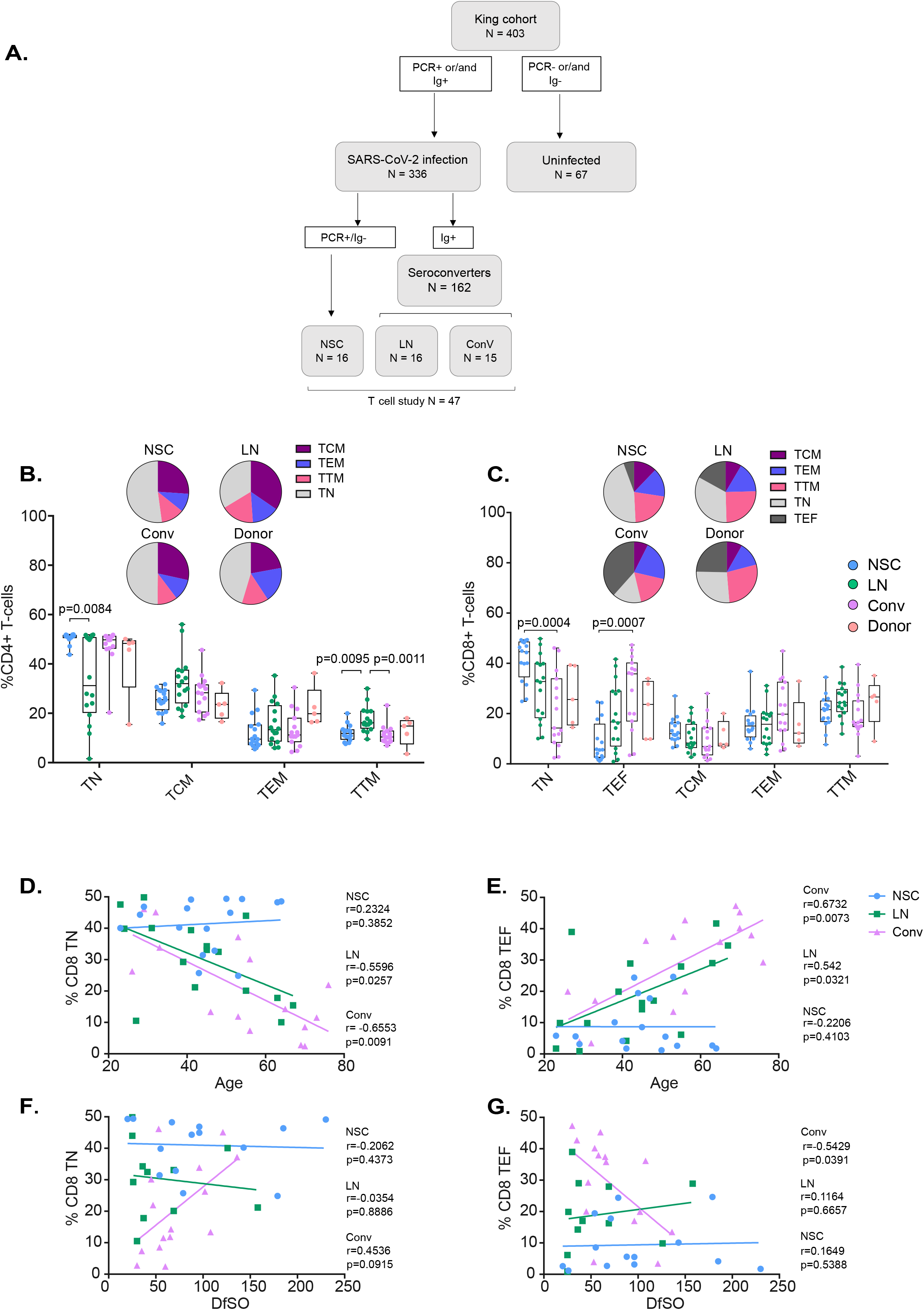
CD4+ and CD8+ T-cell subset distribution in Non-seroconverters. (A) Flow chart of the study groups, including Non-seroconverters (NSC), Low neutralizers (LN), Convalescent (Conv) and blood donor samples (Donor). (B-C) Cryopreserved PBMCs were cultured and surface stained with antibodies targeting CD4 and CD8 as well as T-cell lineage markers CD45RA, CCR7, CD27 to distinguish T-central memory cells (TCM), T effector memory cells (TEM), T transitional memory cells (TTM), T Naïve cells (TN) and T effector cells (TEF). Whisker plots show the frequencies of CD4+ T-cell subsets and pie charts represent the frequency distribution of CD4+ and CD8+ T-cell subsets in each group (Panel B and C, respectively). Statistical analyses were performed using non-parametric ANOVA (Kruskal-Wallis, adjusted for multiple comparisons). Only significant values are indicated in the figure. Linear regression was performed plotting relative frequencies of CD8+ TN (D) and CD8+ TEF (E) against age of the study participants, as well as relative frequencies of CD8+ TN (F) and CD8+ TEF (G) against days from symptom onset (DfSO). Correlation analysis was performed using the Spearman’s rank correlation test; r- and p-values are reported for each group.

**Table 1.** Clinical characteristics of the study groups.

|  | NSC<br>(n=16) | LN<br>(n=16) | Conv<br>(n=15) |
| --- | --- | --- | --- |
| Sex, Female, n (%) | 14 (88) | 11 (69) | 4 (27) |
| Age at diagnosis of<br>SARS-CoV-2 [median, IQR<sup>1</sup>] | 44.5 [38-52] | 43.5 [29-55] | 56 [33-70] |
| Confirmed Positive PCR, n (%) | 16 (100) | 10 (63) | 14 (93) |
| Positive serology, n (%) | 0 (0) | 16 (100) | 15 (100) |
| Days from symptom initiation to 1st sample<br>[median, range]<sup>2</sup> | 84 [20-230] | 36 [0-158] | 58 [25-136] |
| Days from symptoms to 2<sup>nd</sup> sample [median,<br>range] | 92.5 [60-116] |  |  |
| Symptomatic COVID-19, n (%) | 14 (88) | 10 (63) | 15 (100) |
| Mild infection, n (%) | 16 (100) | 15 (93) | 3 (20) |
| Required hospitalization, n (%) | 1 (6) | 2 (13) | 14 (93) |
| Severe presentation of COVID-19, n (%)<sup>3</sup> | 0 (0) | 1 (6) | 12 (80) |
| Co-morbidities, any comorbidity, n (%) | 11 (69) | 8 (50) | 11 (73) |
| HTA | 1 (6) | 1 (6) | 5 (33) |
| Obesity | 0 (0) | 1 (6) | 5 (33) |
| Allergies | 3 (19) | 4 (25) | 3 (20) |
| Asthma | 1 (6) | 0 (0) | 2 (13) |
| Primary immunodeficiency | 1 (6) | 0 (0) | 0 (0) |
| Autoimmunity | 1 (6) | 0 (0) | 0 (0) |
| Neutralizing titer (pseudovirus), reciprocal<br>dilution, median [IQR] | <60<sup>4</sup> | 203 [113-321] | 938 [659-2387] |

NSC: Non-seroconvertors, LN: Low-neutralizers, Conv: Convalescent
<sup>1</sup> [IQR]: inter-quartile range,
<sup>2</sup> Days from PCR to 1st sample if asymptomatic
<sup>3</sup> Severe presentation of SARS-CoV-2: O2 saturation <94%; <sup>4</sup> <60, below the limit of detection

### Determination of anti-SARS-CoV-2 antibodies by Enzyme-linked Immunosorbent Assays

The presence of anti-SARS-CoV-2 antibodies in serum or plasma samples was evaluated using two independent Enzyme-linked Immunosorbent Assays (ELISA). The first was an in-house developed sandwich-ELISA. Briefly, Nunc MaxiSorp ELISA plates were coated overnight at 4°C with 50 ml of capture antibody (anti-6xHis antibody, clone HIS.H8; ThermoFisher Scientific) at 2 mg/mL in PBS. After washing, plates were blocked for two hours at room temperature using PBS containing 1% of bovine serum albumin (BSA, Miltenyi biotech). 50 ml (1mg/mL in blocking buffer) of the following SARS-CoV-2 derived antigens: S1+S2 subunits of the Spike (S) protein and receptor-binding domain (RBD, Sino Biological) were subsequently added and incubated overnight at 4°C. Each plasma sample was evaluated in duplicates at a 1/100 dilution in blocking buffer for each antigen. Antigen free wells were also assessed in parallel for each sample in the same plate to evaluate sample background. Serial dilutions of a positive plasma sample were used as standard. A pool of 10 SARS-CoV-2 negative plasma samples, collected before June 2019, were included as the negative control. Samples were assayed at 1/100 dilution in blocking buffer for one hour at room temperature. The following reagents were used as secondary antibodies: HRP conjugated (Fab)2 Goat anti-human IgG (Fc specific) (1/20000), Goat anti-human IgM (1/10000), and Goat anti-human IgA (alpha chain specific) (1/20000) (all from Jackson Immunoresearch). Secondary antibodies were incubated for 30 minutes at room temperature. After washing, plates were revealed using o-Phenylenediamine dihydrochloride (OPD) (Sigma Aldrich) and the enzymatic reaction was stopped with 4N of H2SO4 (Sigma Aldrich). The signal was analysed as the optical density (OD) at 492 nm with noise correction at 620 nm. The specific signal for each antigen was calculated after subtracting the background signal obtained for each sample in antigen-free wells. The second ELISA was a commercially available IgM and IgG class antibody ELISA against the SARS-CoV-2 NP (ImmunoDiagnostics, Hongkong). Briefly, serum samples were diluted 1:100 and incubated for 1 hour at room temperature. Anti-NP antibodies were captured by immobilized NP recombinant protein. After incubation, captured antibodies were measured by an absorbance microplate reader at 450 nm. The antibody results were expressed as an index value, calculated as the ratio of the OD value for each sample to the OD value of the cut-off (0,200). The test was considered positive when the index value was ≥ 1.1, borderline when index value was ≥ 0.9 to < 1.1, and negative when index value was < 0.9.

### Pseudovirus neutralization assay

HIV reporter pseudoviruses expressing SARS-CoV-2 S protein and Luciferase were generated. pNL4-3.Luc.R-.E- was obtained from the NIH AIDS repository (Connor RI, Chen BK, 195). SARS-CoV-2.SctΔ19 was generated (Geneart) from the full SARS-CoV-2 S gene sequence with a deletion of the last 19 C-terminal codons (Ou et al., 2020), human-codon optimized and inserted into pcDNA3.4-TOPO. Expi293F cells were transfected using the Expifectamine Reagent (Thermo Fisher Scientific, Waltham, MA, USA) with pNL4-3.Luc.R-.E- and SARS-CoV-2.SctΔ19 at an 8:1 ratio, respectively. Control pseudoviruses were obtained by replacing the S protein expression plasmid with a VSV-G protein expression plasmid as reported previously (Ou et al., 2020). Supernatants were harvested 48 h after transfection, filtered at 0.45 µm, frozen and titrated on HEK293T cells overexpressing WT human ACE-2 (Integral Molecular, USA). For the neutralization assay, 200 TCID50 of pseudoviral supernatant was preincubated with serial dilutions of heat-inactivated serum or plasma samples (ranging from 1/60 to 1/14580) for 1h at 37°C and then added to ACE2-overexpressing HEK293T cells. After 48 h, cells were lysed with Britelite Plus Luciferase reagent (Perkin Elmer, Waltham, MA, USA), and luminescence was measured for 0.2 s with the EnSight Multimode Plate Reader (Perkin Elmer). Data were fitted to a four-parameter logistic curve with variable slope using Graph Pad Prism software (v8.3.0). IC50 values are expressed as reciprocal dilution.

### Immunophenotype of S and NP SARS-CoV-2 T-cell responses

To measure SARS-CoV-2 specific T-cell responses, cryopreserved PBMCs were stimulated with total S and NP recombinant proteins (5 μg/mL, Sinobiological, China), Staphylococcal enterotoxin B (SEB) (1 μg/mL, Sigma-Aldrich), or no stimuli, in the presence of CD28/49d co-stimulatory molecules (1 μg/mL, BD) for 17 h at 37 °C in a 5% CO_2_ incubator. After incubation, PBMCs were treated with Monensin A (1 μg/mL, BD Golgi STOP, Thermo Fisher Scientific) for 6h at 37 °C and stored overnight at 4 °C. The next day, cells were stained with a combination of T-cell lineage markers and functional markers for immunophenotyping. In brief, cells were labelled with a viability dye (APC-Cy7, Thermo Fisher Scientific) for 30 min at room temperature (RT) and surface stained for 30 min at RT with anti-human antibodies for CD3 (A700, clone UCHT1, BD), CD4 (FITC,clone OKT4, Biolegend), CD8 (V500, clone RPA-T8, BD), CD45RA (BV786, clone HI100, BD), CCR7 (PE-CF594, clone 150503, BD), CD27 (BV605, clone L128, BD), PD-1 (BV421, clone EH12.1, BD) and activation induced markers (AIM) CD25 (A647, clone BC96, Biolegend), OX40 (PE, clone Ber-ACT35, Biolegend) and CD137 (PeCy7 clone 41BB, Biolegend). Afterwards, cells were fixed with Fix/Perm Buffer A (Thermo Fisher Scientific) for 15 min at RT and stained intracellularly with Fix/Perm Buffer B and antibodies for TNF (PE-Cy7, clone MAb11, BioLegend), IFN-γ (BV711, clone B27, BD), and IL-2 (BV650, clone MQ1-17H12, BD) for 20 min at RT. Finally, cells were resuspended and fixed in formaldehyde 1% and acquired on LSR Fortessa cytometer using FACSDiVa software (BD). Data analysis was performed using FlowJo software version 10.0.7 (Tree Star, Ashland, OR, USA). Specific gates were defined using fluorescence minus one (FMO) controls. Surface markers were measured in total CD4+ and CD8+ T cells. The CD4+ and CD8+ T-cell subsets were differentiated based on CD27 and CCR7 expression: naïve (TN: CD45RA^+^, CD27^+^, CCR7^+^), effector (TEF: CD45RA^+^, CD27^−^, CCR7^−^, only for CD8+), central memory (TCM: CD45RA^−^, CD27^+^, CCR7^+^), transitional memory (TTM: CD45RA^−^, CD27^+^, CCR7^−^), and effector memory (TEM: CD45RA^−^, CD27^−^, CCR7^−^), as previously described (Klatt et al., 2014; Blanch-lombarte et al., 2019) (**Supplementary Fig. S1A-B**). The frequency of T-cell cytokine responders against S and/or NP proteins were defined by the presence of at least one cytokine. In addition, the frequency of total T-cell responders against S and/or NP proteins were defined by the presence of at least one cytokine and/or AIM. We used a 0.2% cut-off value for positivity following background subtraction.

### Statistical analyses

Descriptive and comparison tests were performed using Graph Pad Prism, version 6 (GraphPad Software, Inc., San Diego, CA, USA). Non-parametric ANOVA (Kruskal Wallis) tests were used for groupwise comparisons and adjusted for multiple comparisons. Wilcoxon matched-pairs signed-rank tests were used to compare parameters between sampling time points. Spearman’s rank correlation coefficient test was used for correlation analysis. Non-parametric ANOVA was used to compare differences in age between study groups and was adjusted for multiple comparisons (Dunn’s test). In the analysis of T-cell subsets, median values within phenotypes were normalized to 100%. Single-cell analysis were performed using the statistical package R (v3.6.3). Cells were compensated and selected based on its antigen expression (TNF, IL-2 and IFN-γ intensity) and normalized before performing UMAP dimensionality reduction. We used FlowJo 10.0.7 for the graphical representation of the single-cell analysis.

## RESULTS

### Non-Seroconvertors present skewed T cell distribution towards higher naïve and lower effector CD8+ T cells

Previous studies have demonstrated alterations in immunological parameters following SARS-CoV-2 infection, including lymphopenia with a marked decrease in CD8+ T cells, as well as changes in frequencies of CD8+ T-cell subsets compared to healthy donors (Mathew et al., 2020; Zhao et al., 2020). We determined the differentiation profiles of total T cells in NSC by staining for CD3, CD4 and CD8 markers and differentiation markers (CD45RA/RO, CCR7 and CD27) and compared between study groups (**Fig 1A**, **S1A-B**). No alterations in total frequencies of CD3+ T cells or CD4+/CD8+ ratios were found between SARS-CoV-2 study groups nor when compared with healthy donors (data not shown). By contrast, analysis of CD4+ T-cell subsets revealed higher levels of CD4+ T naïve cells in NSC compared to LN and (p=0.0084, **Fig 1B**) lower levels of T transitional memory cells (TTM) in NSC and Conv compared to LN (p=0.0095 and p=0.0011, respectively, **Fig 1B**). No significant differences were observed regarding the distribution of the other CD4+ T-cell subsets (**Fig 1B**). Interestingly, we observed marked differences in CD8+ T-cell subsets, when comparing NSC and Conv groups (**Fig 1C**). Specifically, NSC had significantly higher frequencies of CD8+ T naïve (TN, median 44.6% vs 14.3%; p=0.0004) and significantly lower frequencies of CD8+ T effector cells (TEF) (median 5.6% vs 35.8%; p=0.0007, Kruskal-Wallis). In NSC, the observed T-cell subset distribution was stable up to a median of 92.5 days of follow-up after infection (**Fig S1C-D**). To assess whether the difference in CD8+ TN and TEF distribution in NSC was related to the individuals’ age or to days from symptom onset (DfSO), we performed correlation analyses. We did not find a significant correlation between age and frequencies of CD8+ TN nor TEF in NSC (**Fig 1D-E**). By contrast, there was a significant negative correlation between frequency of CD8+ TN and age (Spearman r –0.5596, p=0.0257 and r-0.6553, p=0.0091) and a significant positive correlation between frequency of CD8+ TEF and age (r= 0.5420, p= 0.0321 and r=0.6732, p= 0.0073) in the LN and Conv groups, respectively (**Fig 1D-E**). Similarly, while we did not find a significant correlation between DfSO and CD8+ TN or TEF in NSC, a negative correlation between CD8+ TEF % and DfSO in Conv (r= −0.5429, P=0.0391) was observed (**Fig 1F-G**). Taken together, these data indicate a skewed CD8+ T cell subset distribution in NSC towards high levels of naïve and low levels of effector cells independent of sampling time point or patient’s age.

### Highly functional SARS-CoV-2-specific CD4+ and CD8+ T cell responses in Non-Seroconverters

The importance of functional, virus-specific T-cell responses in SARS-CoV-2 disease outcome has been consistently reported (Rydyznski Moderbacher et al., 2020; Liao et al., 2020). Furthermore, T-cell responses against S, NP and M SARS-CoV-2 proteins have been identified in convalescent seronegative individuals (Schulien et al., 2021; Schwarzkopf et al., 2021). To evaluate SARS-CoV-2 specific T cell responses in NSC and the comparative study groups, we stimulated PBMCs with S and NP SARS-CoV-2 proteins and performed intracellular cytokine staining for IFN-γ, IL-2 and TNF production (**Fig 2**). Visualization of cytokine profiles in antigen specific T cells as performed by UMAP analysis is represented in **Fig S2**. Analysis of S and NP-specific CD4+ T-cell responses revealed antigen-specific cytokine production in all study groups (**Fig 2A)** and no significant differences in the frequency of S and NP-specific CD4+ T cells between NSC and the other SARS-CoV-2 groups for any of the cytokines studied. However, LN displayed significantly lower NP-specific IFN-γ production than Conv (p=0.0402). Lack of statistically significant differences between NSC and Conv might be associated with the high interindividual variability, despite the lower median frequency of NP-specific CD4+ T cell responses in NSC vs. Conv individuals. The difference in median frequency was particularly marked for the proinflammatory cytokine TNF in NSC compared to Conv (0.07% vs 1% in TNF, 0.01% vs 0.3% in IFN-γ and 0% vs 0.26% in IL-2, respectively; **Fig 2A**). Similarly to SARS-CoV-2-specific CD4+ T cells, S and NP-specific CD8+ T-cell responses were present in all study groups. No statistically significant differences were found between them and marked interindividual variability was observed (**Fig 2B**). Also, median frequencies of IFN-γ or TNF producing NP-specific CD8+ T cells were lower in NSC vs Conv (0.06% vs 0.23% in TNF and 0.05% vs 0.24% for IFN-γ, respectively; **Fig 2B**). Next, we determined the frequency of T-cell cytokine responders across study groups defined by the presence of either S or NP CD4+ or CD8+ T cells producing at least one cytokine and using a threshold of 0.2% cut-off for a positive response after background subtraction. Overall, the frequency of SARS-CoV-2 responders were 81% in NSCs, 75% in LN, 80% in Conv, and 20% of healthy donors. Distributed by SARS-CoV-2 proteins, we observed a frequency of S responders in 56% of NSCs, 50% of LN, 53% of Conv, and 20% of healthy donors. The frequency of NP responders was 50% of NSCs, 44% of LN, 73% of Conv, and 0% of healthy donors. These data are consistent with previous studies demonstrating the presence of SARS-CoV-2 reactive T cells in seronegative SARS-CoV-2 convalescent individuals and in up to 50% of unexposed individuals (Grifoni et al., 2020; Mateus et al., 2020; Sekine et al., 2020).

**Figure 2.**
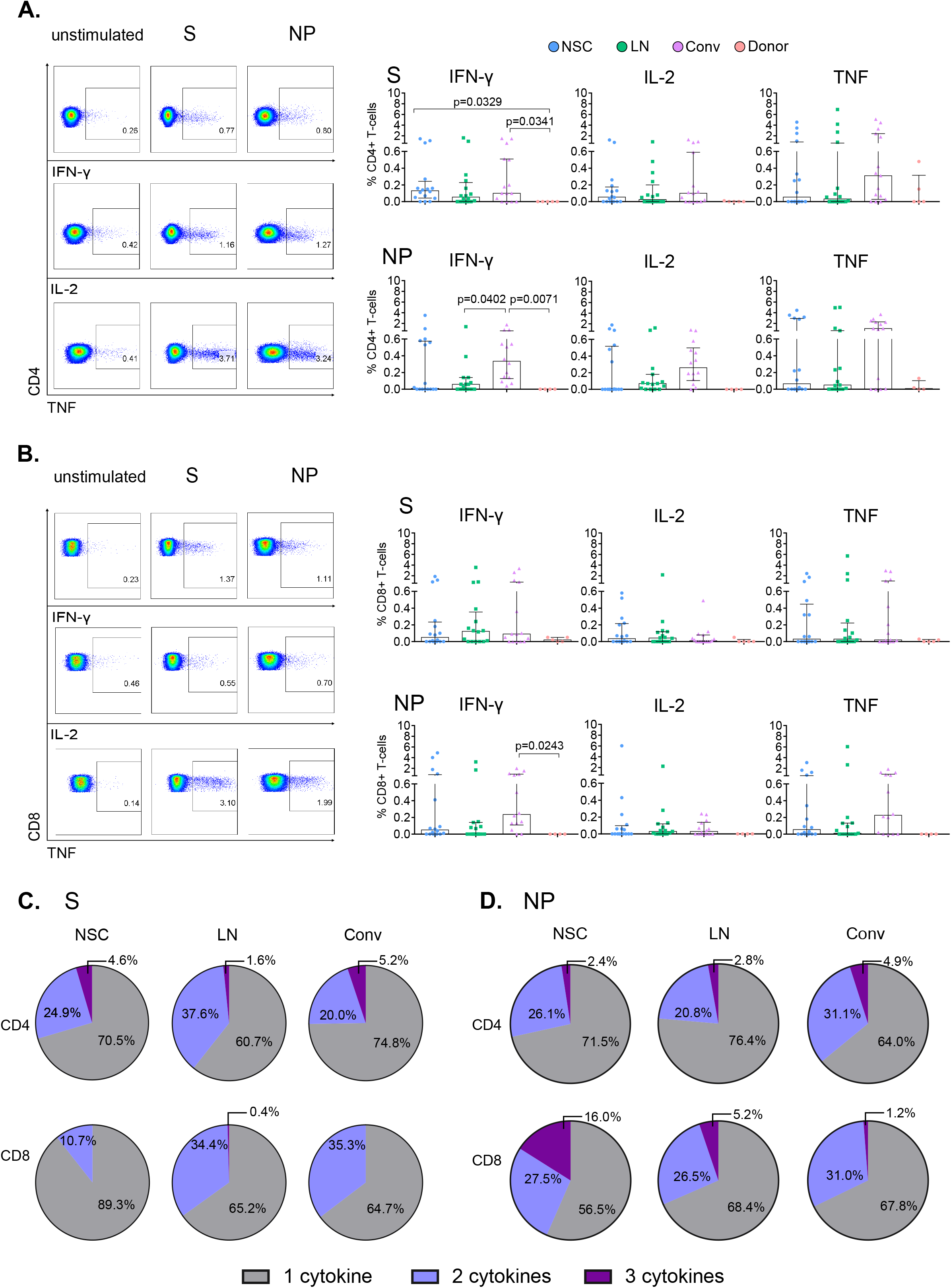
T cells of SARS-CoV-2 Non-seroconverters produce (polyfunctional) pro-inflammatory cytokines in response to Spike and Nucleocapsid. Cryopreserved PBMCs from SARS-CoV-2 Non-seroconverters (NSC) and comparative study groups, Low neutralizers (LN), Convalescent (Conv), and healthy blood donor samples (Donors) were stimulated for 17 hours with Spike (S) and Nucleocapsid proteins (NP). An unstimulated condition was used as control. Production of IFN-γ, IL-2, and TNF was monitored by Flow cytometric analysis following intracellular cytokine staining. (A-B) Left panels depict the gating strategy for analysis of cytokine expression and right panels represent the frequencies of IFN-γ, IL-2, and TNF in CD4+ (A) and CD8 (B) T cells in all study groups in response to S and NP. Pie charts demonstrate the relative proportions of T cells producing one, two or three cytokines in response to S in CD4+ T cells and CD8+ T cells (C). Relative proportions of T cells producing one, two or three cytokines in response to NP in CD4+ T cells and CD8+ T cells (D). Statistical analysis was performed using Non-parametric ANOVA (Kruskal Wallis) adjusted for multiple comparisons. Error bars represent interquartile range and median values per group are shown. Only significant differences are indicated in the figure.

Polyfunctional T cells have been associated with protective antiviral immune responses (Seder et al., 2008). Lower frequencies of T helper type 1 cells have been reported in hospitalized COVID-19 patients when compared to influenza-reactive T cells (Meckiff et al., 2020) and decreased frequencies of polyfunctional CD4+ T cells have been associated with severe SARS-CoV-2 infection. In addition, elevated frequencies of non-functional T cells, defined by the absence of IFN-γ, TNF and IL-2 production have been related to severe SARS-CoV-2 disease course (Zheng et al., 2020). We next assessed the polyfunctionality of S and NP-specific T cells in NSCs and comparative study groups. Polyfunctionality was assessed by Boolean gating using IFN-γ, IL-2 and TNF in CD4+ and CD8+ T cells. In response to S proteins, CD4+ T cells of NSCs displayed similar characteristics as did Conv (with 4.6% vs 5.2% trifunctional cells, and 24.9% vs 20% bifunctional cells respectively), while LN had lower frequencies of trifunctional cells (1.6%), but higher frequencies of bifunctional cells (37.6%; **Fig 2C**). On the other hand, CD8+ T cells in response to S displayed higher frequencies of monofunctional cells in NSC (89.3%) compared with LN (65.2%) and Conv (64.7%), and lower frequencies of bifunctional cells (10.7%) compared to LN (34.4%) and Conv (35.3%). Negligible frequencies in CD8+ trifunctional cells were observed in all groups (**Fig 2C**). In response to NP proteins, CD4+ T cells of NSCs displayed similar characteristics as did Conv (with 2.4% vs 4.9% trifunctional cells, and 26.1% vs 31.1% bifunctional cells respectively) (**Fig 2D**). Interestingly, NSC had the highest frequencies of trifunctional NP-specific CD8+ T cells, accounting for 16% of their NP-specific CD8+ T cells, compared with LN and Conv (5.2% and 1.2%, respectively, **Fig 2D**). By contrast, no major differences in mono and bifunctional CD8+ T cells between groups were observed (**Fig 2D**). These results demonstrate a particularly elevated frequency of highly functional SARS-CoV2-specific T cells responses, especially polyfunctional NP-specific CD8+ T cell responses as a hallmark of NSC.

### Non-seroconverters display lower levels of T-cell activation-induced markers and PD-1

Next, we used PD-1 and activation-induced markers (AIM) to evaluate total T cell activation and activation status of SARS-CoV-2 specific T cells (Sekine et al., 2020; Grifoni et al., 2020). AIMs have been used to identify antigen-specific CD4+ T cells via the detection of upregulated surface markers following antigen stimulation (Reiss et al., 2017). We assessed general T-cell activation and AIMs using antibodies directed against CD25, OX40 and CD137 and PD-1 in unstimulated and S- and NP-antigen stimulated conditions (**Fig 3A-C**). As observed for SARS-CoV-2-specific T-cell cytokine production, a high level of interindividual variability was observed within groups for AIMs. Still, we found statistically significantly lower expression of CD25^+^OX40^+^ CD4^+^ T cells in response to S in NSC compared to Conv (median 0.13% vs 0.59%, respectively, p=0.0216, **Fig 3B**). The same was observed for S-specific CD4+ CD137^+^OX40^+^ T cells (p=0.0117, data not shown). In addition, CD25^+^OX40^+^ CD4+ T cells in response to NP were not significantly different between NSC and the rest of the groups, although lower median frequencies were observed (0.005% NSC vs 0.84% Conv, **Fig 3B**). The analysis of AIM on CD8+ T cells did not reveal significant differences in CD137^+^ CD8^+^ T-cell frequencies between study groups (**Fig 3B)**. However, the median frequency of CD137^+^ CD8^+^ T cells was lower in NSC, particularly in response to NP (0% NSC vs 0.77% Conv, **Fig 3B**). Also, we performed longitudinal follow-up of AIM expression in samples from 6 NSC individuals. Frequencies of S and NP CD25^+^OX40^+^ CD4^+^ T cells decreased or remained undetectable between sampling time points in all the NSC except for one individual, who also displayed increased cytokine production in response to S (**Fig S1E**). A comparable profile was observed for S and NP CD137+ CD8+ T cells over time (**Fig S1E**).

**Figure 3.**
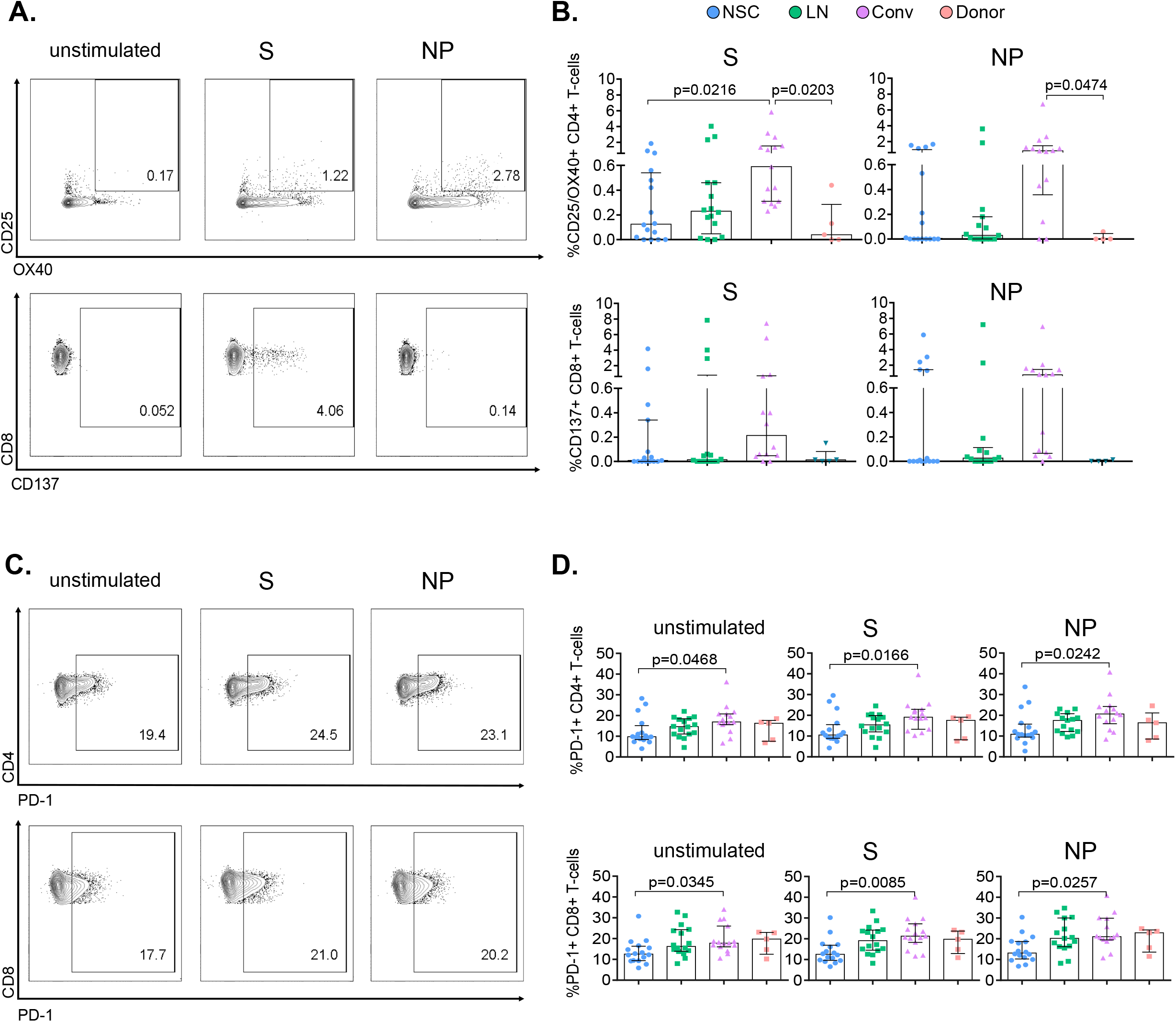
Expression of activation markers is lower in SARS-CoV-2 Non-seroconvertors compared to seropositive convalescent individuals. PBMCs from Non-seroconvertors and comparative study groups were stimulated for 17 hours with S and NP recombinant proteins and stained using antibodies directed against activation-induced markers (AIM; CD25, OX40, CD137) and PD-1, and analysed by flow cytometric analysis. (A) Gating strategy for AIM expression in CD4+ and CD8+ T cells. (B) Dot plots showing expression of activation induced markers in CD4+ (CD25^+^OX40^+^) and CD8+ T cells in response to S and NP (CD137^+^). Values from unstimulated cells were subtracted for each individual. (C) Gating strategy for analysis of PD-1 expression on CD4+ and CD8+ T cells. (D) Expression of PD-1 on CD4+ and CD8+ T cells in unstimulated cells as well as following S and NP stimulation. Error bars represent interquartile range and median values per group are shown. Statistical analysis was performed using Kruskal-Wallis non-parametric ANOVA adjusted for multiple comparisons, only significant differences are highlighted in the figure.

Along with decreased functionality, elevated activation and exhaustion markers such as PD-1 in T cells from patients experiencing severe COVID-19 have been described (Zheng et al., 2020). Analysis of PD-1 revealed a significant reduction in the frequency of PD-1 expressing T cells in NSC compared to convalescent individuals in all conditions tested (**Fig 3D**) in CD4+ (unstimulated, p=0.0468; S, p=0.0166 and NP, p=0.0242) and CD8+ T cells (unstimulated, p=0.0345; S, p=0.0085, and NP, p=0.0257). Longitudinal follow-up of PD-1 expression in NSC demonstrated stable frequencies of PD-1 expression in CD4+ and CD8+ T cells independent of S or NP antigen stimulation (**Fig S1F**). Only a decrease in the frequency of PD-1^+^ CD4^+^ T cells following S-stimulation was observed over time (p=0.0313, Wilcoxon matched-paired signed-rank test, **Fig S1F**). Overall, these data indicate lower levels of AIM expression (CD25^+^OX40^+^) in CD4+ T cells in response to S and an overall low level of PD-1 expression in CD4+ and CD8+ T cells as particular immunological traits of NSC.

## DISCUSSION

While unprecedented advances have been made for the approval of COVID-19 vaccines with a high degree of efficacy (Logunov et al., 2021; Voysey et al., 2021; Mahase, 2020; Oliver et al., 2020), ensuring long-lasting immune protection remains a potential challenge for current vaccines. Severe COVID-19 has been related to increased age, high levels of inflammatory markers and seroconversion (Staines et al., 2021). Persistent seroconversion and higher neutralizing titers have been found consistently in hospitalized SARS-CoV-2 infected individuals (Trinité et al., 2021; Pradenas et al., 2021). These studies support the relevance of alternative immune mechanisms to humoral responses crucial in limiting SARS-CoV-2 pathogenesis. In this study, we have conducted a detailed immune characterization of T-cell responses in SARS-CoV-2 individuals, suggesting that virus-specific T-cell responses in the absence of seroconversion may mediate immune protection from disease severity and viral pathogenesis.

Non-Seroconvertors (NSC) in SARS-CoV-2 infection have been described to be present at a frequency of 2-17% among convalescent individuals (Schulien et al., 2021; Schwarzkopf et al., 2021; Sekine et al., 2020). While previous studies have been shown the presence of SARS-CoV-2 specific T-cell responses in NSC (Schwarzkopf et al., 2021; Sekine et al., 2020), and SARS-CoV-2 CD8+ T-cell responses have been very well characterized in a small fraction of those (Schulien et al., 2021), a thorough characterization of the NSC immune phenotype is still missing. Here, we overcome some of the previous studies limitations through a detailed identification of NSC and comparative analyses of total and S and NP-specific T cells. These data are essential to strengthen our understanding of T-cell mediated immunity and answer important questions regarding the identification of distinctive immunological traits associated with immune protection against SARS-CoV-2.

We identified 4.8% of Non-seroconvertors in the KING extension cohort. We used a conservative approach to identify NSC by including confirmed SARS-CoV-2 PCR positivity, double ELISA testing and sampling-time estimation to allow for seroconversion. Longitudinal follow-up in the absence of SARS-CoV-2 IgG, IgM and IgA seroconversion over time further confirm the true nature of NSC in our study. Epidemiologically, NSCs were biased towards individuals with mild to moderate SARS-CoV-2 infection and a higher frequency of females. These data contrast with the presence of severe infection in 80% of Conv and higher frequency of males. The observed sex distribution is consistent with previous findings suggesting females exhibit robust T-cell responses and are less prone to severe SARS-CoV-2 infection and death than males (Takahashi et al., 2020; Jin et al., 2020; Vahidy et al., 2021). No significant differences in age distribution between NSC and Conv were found excluding age as one of the factors accounting for the differences in disease severity observed.

We performed a characterization of T-cell immunity in NSC and compared with SARS-CoV-2 infected individuals stratified by seroconversion, magnitude of humoral neutralizing activity, and disease severity. In terms of T-cell immunophenotype, we observed a skewed distribution of CD8+ T cell subsets with a higher frequency of naïve and lower frequency of effector CD8+ T cells in NSC. The observed distribution was independent of age and DfSO. These data contrast with the negative correlation between the frequency of naïve CD8+ T cells and the positive correlation between the frequency of effector and CD8+ T cells with age found in Conv and LN. Of note, this is in line with previous observations indicating a decline in the frequencies of naïve T cells with age and the association of low levels of naïve CD8+ with increased risk of severity in COVID-19 (Hong et al., 2004)(Rydyznski Moderbacher et al., 2020). The skewed CD8+ T cell distribution independent of age and time observed support a specific characteristic of T cell homeostasis in NSC. The higher levels of CD8+ TN cells in NSC could be the result of a continuous thymic repopulation of the naïve compartment, as observed in HIV-1 viremic non-progressor individuals (Singh et al., 2020). High levels of CD8+ TN could also allow rapid and efficient priming and expansion of SARS-CoV-2-specific CD8+ T cells favoring viral control. The low levels of CD8+ TEF observed in NSC could partially be explained by highly functional CD8+ T cells, overcoming the need for a large proportion of effector cells.

We extended the characterization of T-cell immunity to functional SARS-CoV-2-specific CD4+ and CD8+ T-cell responses by cytokine production and AIMs in response to S and NP proteins. Overall, we did not observe differences in cytokine production in CD4+ and CD8+ T cells in response to S and NP proteins in NSC and the rest of the groups, which may have been due to the marked interindividual variability in these parameters. This is in partial agreement with data reporting the targeting of T-cell responses toward S-, M- and N-proteins in COVID-19 patient samples to be mainly uniform across disease severities (Thieme et al., 2020). However, in NSC S and NP-specific T cells appear to have a balanced profile of IFN-γ, IL-2 and TNF production, while Conv exhibits higher median frequencies of TNF producing cells. TNF secretion by CD4+ T cells serves as a costimulatory signal for B cell activation (Aversa et al., 1993), which is in line with the high levels of seroconversion and plasma neutralizing activity reported in convalescent hospitalized individuals. Moreover, elevated levels of TNF may be both beneficial and detrimental. While TNF is one of the main pro-inflammatory cytokines produced by T cells responding to viral, bacterial, and fungal infections (Rahman and McFadden, 2006), its effects are also implicated in increased viral replication (Kumar et al., 2013; Roca et al., 2008; Espín-Palazón et al., 2016). Although NSC displayed tendencies towards lower median frequencies of SARS-CoV-2-specific T cell responses, we measured high levels of IFN-γ^+^ IL-2^+^ TNF^+^ NP-specific CD8+ T cells in NSC. The presence of polyfunctional NP-specific CD8+ T cells in NSC emphasized the importance to understand quality over quantity in the context of immune protection by T cell responses in SARS-CoV-2 infection. These findings follow previous studies supporting a distinctive contribution of surface and non-surface antigen- specific-CD8+ T cells to disease control and the potential role of NP-specific and other non-S CD8+ T-cell responses in immune protection (Peng et al., 2020). Indeed, CD8+ T-cell responses were shown to be more likely to target NP than S (Cohen et al., 2021). Our data is also in line with previous findings showing NP-specific CD8+ T cells directed against the immunodominant B7/N105 epitope were detected at high frequency in pre-COVID-19 samples, and even more so during acute COVID-19 and convalescence. A predominantly naïve phenotype with high TCR plasticity was observed among these cells (Nguyen et al., 2021).

Complementary to cytokine producing cells, we investigated the expression of AIMs in CD4+ T cells (CD25^+^OX40^+^) and CD8+ T cells (CD137^+^) and PD-1 in response to S and NP antigens. Data from AIMs identified lower median frequencies of S CD25^+^OX40^+^ CD4+ T cells and a trend towards lower NP CD137^+^ CD8^+^ T cells as distinctive traits in NSC. Higher frequencies of AIM^+^ T cells observed in Conv are likely to detect alternative cytokine profiles and T regulatory cells as described previously (Reiss et al., 2017). Another explanation could be an association between the pro-inflammatory environment in Conv and the overall increase in immune activation found.

These data highlight the importance of the quality of SARS-CoV-2 T-cell responses, as the NSC displayed polyfunctional T cells despite lower overall expression of AIMs. These findings have additional value considering the worldwide expansion of SARS-CoV-2 variants of concern for vaccine efficacy (Kirby, 2021; Tegally et al., 2021). In this context, the quality of antigen-specific CD4+ and CD8+ T-cell responses may be of crucial importance to recognize emerging SARS-CoV-2 variants.

Total frequencies of SARS-CoV-2 T-cell responders, taking into account both cytokines and AIMs were 81% of NSCs, 69% of LN, 100% of Conv, and 20% of healthy donors responded to either S or NP consistent with previous studies (Sekine et al., 2020; Mateus et al., 2020). We did not find T-cell responses in 19% of NSCs to S nor NP proteins. These data support intrinsic diversity within the NSC phenotype in terms of other cell types or T cells targeting alternative regions of the proteome not included in our analyses.

Moreover, we observed decreased levels of PD-1 in CD4+ and CD8+ T cells in NSC compared to Conv with and without SARS-CoV-2 antigen stimulation. Elevated activation and exhaustion markers have been found in T cells of severe COVID-19 patients (Zheng et al., 2020). Aside from elevated activation and exhaustion markers such as PD-1, decreased functionality has been observed in T cells from severe COVID-19 patients (Zheng et al., 2020). Recently, SARS-CoV-2 specific CD8+ T cells expressing PD-1 were shown to be functional rather than exhausted, as they also produced IFN-γ (Rha et al., 2021). Therefore, while the PD-1 expressing cells in Conv may be functional, our data suggest that T cells of NSC can exhibit functionality despite reduced PD-1 and AIMs expression levels. This is potentially explained by pre-existing cross-reactive T-cell responses.

Given the reports on pre-existing immunity in uninfected individuals, it is tempting to speculate that pre-existing T cell immunity derived from previous exposure to other coronaviruses mediate the NSC phenotype. This would be in line with the generation of a rapid response without the need for high levels of activation, it is also a plausible explanation of why the NSC can control SARS-CoV-2 with T cells in the absence of antibodies. Indeed, cross-reactive memory CD4+ T cells with comparable affinity to epitopes from SARS-CoV-2 and common cold coronaviruses have been described in unexposed individuals with the most frequently targeted region being S (Mateus et al., 2020). Reactivity to NP has previously been demonstrated in 30% of SARS-CoV-1 and SARS-CoV-2 unexposed individuals. A mapped epitope at aa 101-120 shared a high degree of homology to the sequences of the NP of MERS-CoV, OC43 and HKU1 (Le Bert et al., 2020).

Our study has some limitations. First, we found a bias in epidemiological characteristics in NSC toward a high frequency of females and median lower age. Second, we used S and NP recombinant proteins as opposed to optimal peptides for the study of antigen-specific T cells, potentially underestimating the true frequency of virus-specific T cells. Third, our functional study of T cells is limited to Th1-type cytokines and does not necessarily reflect the full functionality of antigen-specific T cells (Ruiz-Riol et al., 2015). Further immunological studies of NSC are required for additional information on this phenotype.

Collectively, our data suggest a protective role of highly functional SARS-CoV-2 specific T-cell responses in the absence of humoral responses against severe COVID-19 while maintaining a low degree of T-cell activation. Our study provides important information required for successfully guiding clinical monitoring and designing future immune interventions.

## Acknowledgments

We are deeply grateful to all participants and to the technical staff of IrsiCaixa for sample processing (L. Ruiz, E. Grau, R. Ayen, L. Gomez, C. Ramirez, M. Martinez, T Puig) and staff from the Fight AIDS Foundation (R Toledo, A. Chamorro, J Puig). This study was supported in part by grants from National Health Institute Carlos III (ISCIII) COV20/00660, PI17/000164 and RETIC RD16/0025/0041 (Co-funded by European Regional Development Fund/European Social Fund) for JGP. The funders had no role in study design, data collection and analysis, the decision to publish or drafting of the manuscript. OBL was supported by the grant for Catalan Government and the European Social Fund AGAUR-FI_B 00582 Ph.D. fellowship. This study has received partial funding from Grifols and the the crowdfunding initiatives “https://www.yomecorono.com”, BonPreu/Esclat and Correos. We thank “CERCA Programme/Generalitat de Catalunya for institutional support and the Foundation Dormeur for financial support.

## Author contribution statement

Conceptualization, JGP, AK, MM, BC, CB, JC, JB; methodology EJM, AK, OBL, DO, JC, MM, EMC, BQS, IB, RP; validation, MM, JC, JB, RP, LM, EMC, BQS, IB, AK, JGP; Resources, JGP, BC, MM, JC, JB, RP, LM, ACH; formal analysis; EJM, AK, OBL, DO; investigation, AK, JGP; data curation, MM, JB, JC, EMV, BQS, EJM, AK, OBL, DO; software, DO; visualization, AK, OBL, DO, JGP; writing – original draft preparation AK, JGP; supervision, JGP; writing – review and editing AK, JGP, EJM, EMC, JB, MM, OBL, JC, BQS, CB, DO; project administration, JGP; funding acquisition JGP, BC.

---

**Figure S1.**
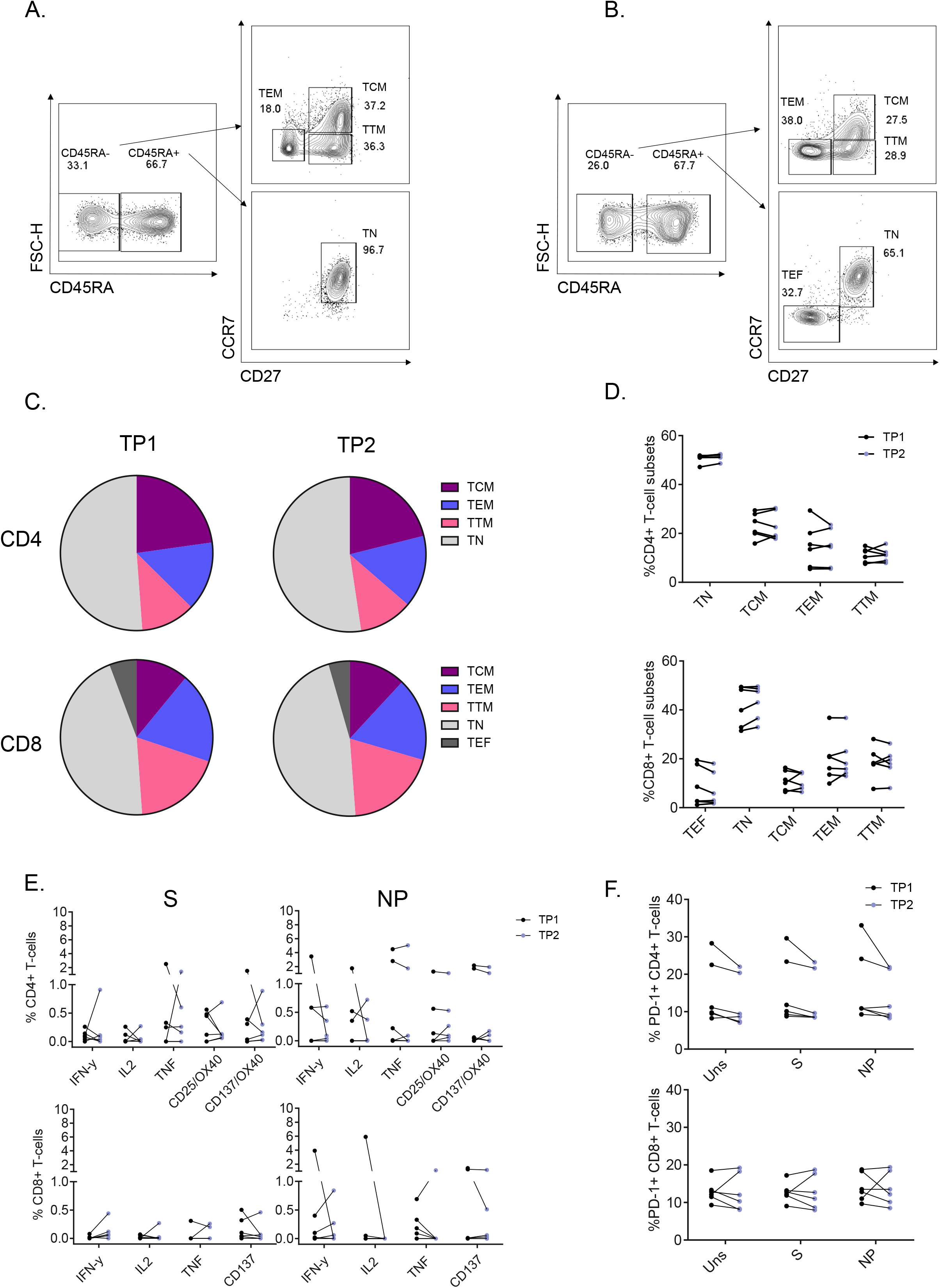
Longitudinal evolution of T-cell subsets and antigen specific responses in SARS-CoV-2 Non-seroconverters. T-cell markers were assessed in two sampling timepoints in Non-seroconvertors. (A) Gating strategy for differentiation of subsets in CD4+ T cells, (B) and CD8+ T cells. Changes in T-cell subset distributions are represented by pie charts showing CD4+ and CD8+ T-cell subset distribution at sampling timepoint 1 and 2 (TP1 and TP2) (C) and graphs representing the normalized subset values within phenotypes (D). Differences in Cytokine production and activation induced markers between S1 and S2 in response to S and NP are shown in (E), and PD-1 expression in CD4+ and CD8+ T cells in unstimulated, S- and NP-stimulated cells at TP1 and TP2 are shown in (F).

**Figure S2.**
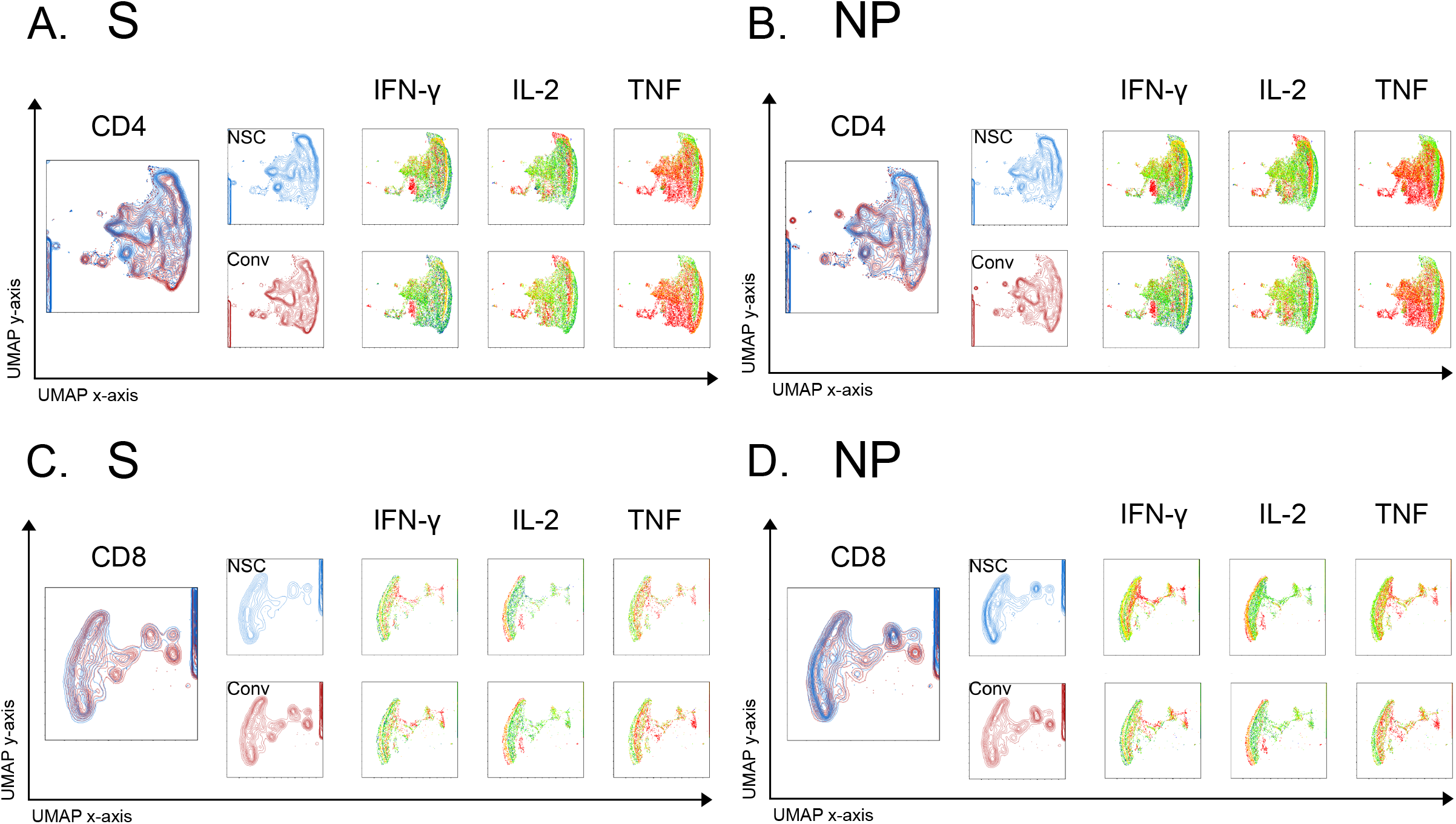
Cytokine expression in CD4+ and CD8+ T cells of Non-seroconvertors and seropositive convalescent individuals. Single-cell analysis based on antigen expression (TNF, IL-2 and IFN-γ intensity) using UMAP dimensionality reduction in CD4+ T-cells following stimulation with S (A) and NP (B) as well as CD8+ T cells stimulated with S (C) and NP (D).

## REFERENCES

Aversa, G., J. Punnonen, and J.E. De Vries. 1993. The 26-kD transmembrane form of tumor necrosis factor α on activated CD4 + T cell clones provides a costimulatory signal for human B cell activation. J. Exp. Med. 177:1575–1585. doi:10.1084/jem.177.6.1575.

Le Bert, N., A.T. Tan, K. Kunasegaran, C.Y.L. Tham, M. Hafezi, A. Chia, M.H.Y. Chng, M. Lin, N. Tan, M. Linster, W.N. Chia, M.I.C. Chen, L.F. Wang, E.E. Ooi, S. Kalimuddin, P.A. Tambyah, J.G.H. Low, Y.J. Tan, and A. Bertoletti. 2020. SARS-CoV-2-specific T cell immunity in cases of COVID-19 and SARS, and uninfected controls. Nature. 584:457–462. doi:10.1038/s41586-020-2550-z.

Blanch-lombarte, O., G. Cristina, B. Revollo, E. Jim, B. Clotet, J.G. Prado, and J. Martinez-picado. 2019. Enhancement of Antiviral CD8 + T-Cell Responses and Complete Remission of Metastatic Melanoma in an HIV-1-Infected Subject Treated with Pembrolizumab. J. Clin. Med. 8:1–11.

Blanco-Melo, D., B.E. Nilsson-Payant, W.C. Liu, S. Uhl, D. Hoagland, R. Møller, T.X. Jordan, K. Oishi, M. Panis, D. Sachs, T.T. Wang, R.E. Schwartz, J.K. Lim, R.A. Albrecht, and B.R. tenOever. 2020. Imbalanced Host Response to SARS-CoV-2 Drives Development of COVID-19. Cell. 181:1036–1045.e9. doi:10.1016/j.cell.2020.04.026.

Breton, G., P. Mendoza, T. Hägglöf, T.Y. Oliveira, D. Schaefer-Babajew, C. Gaebler, M. Turroja, A. Hurley, M. Caskey, and M.C. Nussenzweig. 2021. Persistent cellular immunity to SARS-CoV-2 infection. J. Exp. Med. 218. doi:10.1084/JEM.20202515.

Chang, A.Y., M.R. Cullen, R.A. Harrington, and M. Barry. 2020. The impact of novel coronavirus COVID-19 on noncommunicable disease patients and health systems: a review. J. Intern. Med. 289. doi:10.1111/joim.13184.

Cohen, K.W., S.L. Linderman, Z. Moodie, J. Czartoski, G. Mantus, C. Norwood, L.E. Nyhoff, V. Edara, K. Floyd, S.C. De Rosa, H. Ahmed, R. Whaley, S.N. Patel, B. Prigmore, M.P. Lemos, S. Furth, M.P. Gharpure, S. Gunisetty, A. Stephens, R. Antia, V.I. Zarnitsyna, D.S. Stephens, N. Rouphael, E.J. Anderson, A.K. Mehta, M.S. Suthar, R. Ahmed, and M. Juliana McElrath. 2021. Longitudinal analysis shows durable and broad immune memory after SARS-1 CoV-2 infection with persisting antibody responses and memory B and T cells 2. medRxiv. 2021.04.19.21255739. doi:10.1101/2021.04.19.21255739.

Dan, J.M., J. Mateus, Y. Kato, K.M. Hastie, E.D. Yu, C.E. Faliti, A. Grifoni, S.I. Ramirez, S. Haupt, A. Frazier, C. Nakao, V. Rayaprolu, S.A. Rawlings, B. Peters, F. Krammer, V. Simon, E.O. Saphire, D.M. Smith, D. Weiskopf, A. Sette, and S. Crotty. 2021. Immunological memory to SARS-CoV-2 assessed for up to 8 months after infection. Science (80-.). 371. doi:10.1126/science.abf4063.

Espín-Palazón, R., A. Martínez-López, F.J. Roca, A. López-Muñoz, S.D. Tyrkalska, S. Candel, D. García-Moreno, A. Falco, J. Meseguer, A. Estepa, and V. Mulero. 2016. TNFα Impairs Rhabdoviral Clearance by Inhibiting the Host Autophagic Antiviral Response. PLOS Pathog. 12:e1005699. doi:10.1371/journal.ppat.1005699.

Garcia-Beltran, W.F., E.C. Lam, M.G. Astudillo, D. Yang, T.E. Miller, J. Feldman, B.M. Hauser, T.M. Caradonna, K.L. Clayton, A.D. Nitido, M.R. Murali, G. Alter, R.C. Charles, A. Dighe, J.A. Branda, J.K. Lennerz, D. Lingwood, A.G. Schmidt, A.J. Iafrate, and A.B. Balazs. 2021. COVID-19-neutralizing antibodies predict disease severity and survival. Cell. 184:476–488.e11. doi:10.1016/j.cell.2020.12.015.

Grifoni, A., D. Weiskopf, S.I. Ramirez, J. Mateus, J.M. Dan, C.R. Moderbacher, S.A. Rawlings, A. Sutherland, L. Premkumar, R.S. Jadi, D. Marrama, A.M. de Silva, A. Frazier, A.F. Carlin, J.A. Greenbaum, B. Peters, F. Krammer, D.M. Smith, S. Crotty, and A. Sette. 2020. Targets of T Cell Responses to SARS-CoV-2 Coronavirus in Humans with COVID-19 Disease and Unexposed Individuals. Cell. 181:1489–1501.e15. doi:10.1016/j.cell.2020.05.015.

Hong, M.S., J.M. Dan, J.Y. Choi, and I. Kang. 2004. Age-associated changes in the frequency of naïve, memory and effector CD8+ T cells. Mech. Ageing Dev. 125:615–618. doi:10.1016/j.mad.2004.07.001.

Jin, J.-M., P. Bai, W. He, F. Wu, X.-F. Liu, D.-M. Han, S. Liu, and J.-K. Yang. 2020. Gender Differences in Patients With COVID-19: Focus on Severity and Mortality. Front. Public Heal. 8:152. doi:10.3389/fpubh.2020.00152.

Kiepiela, P., K. Ngumbela, C. Thobakgale, D. Ramduth, I. Honeyborne, E. Moodley, S. Reddy, C. De Pierres, Z. Mncube, N. Mkhwanazi, K. Bishop, M. Van Der Stok, K. Nair, N. Khan, H. Crawford, R. Payne, A. Leslie, J. Prado, A. Prendergast, J. Frater, N. McCarthy, C. Brander, G.H. Learn, D. Nickle, C. Rousseau, H. Coovadia, J.I. Mullins, D. Heckerman, B.D. Walker, and P. Goulder. 2007. CD8+ T-cell responses to different HIV proteins have discordant associations with viral load. Nat. Med. 13:46–53. doi:10.1038/nm1520.

Kirby, T. 2021. New variant of SARS-CoV-2 in UK causes surge of COVID-19. Lancet Respir. Med. 9:e20–e21. doi:10.1016/s2213-2600(21)00005-9.

Klatt, N.R., S.E. Bosinger, M. Peck, L.E. Richert-Spuhler, A. Heigele, J.P. Gile, N. Patel, J. Taaffe, B. Julg, D. Camerini, C. Torti, J.N. Martin, S.G. Deeks, E. Sinclair, F.M. Hecht, M.M. Lederman, M. Paiardini, F. Kirchhoff, J.M. Brenchley, P.W. Hunt, and G. Silvestri. 2014. Limited HIV Infection of Central Memory and Stem Cell Memory CD4+ T Cells Is Associated with Lack of Progression in Viremic Individuals. PLoS Pathog. 10. doi:10.1371/journal.ppat.1004345.

Kumar, A., W. Abbas, and G. Herbein. 2013. TNF and TNF receptor superfamily members in HIV infection: New cellular targets for therapy? Mediators Inflamm. 2013. doi:10.1155/2013/484378.

Kuri-Cervantes, L., M.B. Pampena, W. Meng, A.M. Rosenfeld, C.A.G. Ittner, A.R. Weisman, R.S. Agyekum, D. Mathew, A.E. Baxter, L.A. Vella, O. Kuthuru, S.A. Apostolidis, L. Bershaw, J. Dougherty, A.R. Greenplate, A. Pattekar, J. Kim, N. Han, S. Gouma, M.E. Weirick, C.P. Arevalo, M.J. Bolton, E.C. Goodwin, E.M. Anderson, S.E. Hensley, T.K. Jones, N.S. Mangalmurti, E.T. Luning Prak, E.J. Wherry, N.J. Meyer, and M.R. Betts. 2020. Comprehensive mapping of immune perturbations associated with severe COVID-19. Sci. Immunol. 5. doi:10.1126/sciimmunol.abd7114.

Li, T., J. Xie, Y. He, H. Fan, L. Baril, Z. Qiu, Y. Han, W. Xu, W. Zhang, H. You, Y. Zuo, Q. Fang, J. Yu, Z. Chen, and L. Zhang. 2006. Long-term persistence of robust antibody and cytotoxic T cell responses in recovered patients infected with SARS coronavirus. PLoS One. 1. doi:10.1371/journal.pone.0000024.

Liao, M., Y. Liu, J. Yuan, Y. Wen, G. Xu, J. Zhao, L. Cheng, J. Li, X. Wang, F. Wang, L. Liu, I. Amit, S. Zhang, and Z. Zhang. 2020. Single-cell landscape of bronchoalveolar immune cells in patients with COVID-19. Nat. Med. 26:842–844. doi:10.1038/s41591-020-0901-9.

Logunov, D.Y., I. V Dolzhikova, D. V Shcheblyakov, A.I. Tukhvatulin, O. V Zubkova, A.S. Dzharullaeva, A. V Kovyrshina, N.L. Lubenets, D.M. Grousova, A.S. Erokhova, A.G. Botikov, F.M. Izhaeva, O. Popova, T.A. Ozharovskaya, I.B. Esmagambetov, I.A. Favorskaya, D.I. Zrelkin, D. V Voronina, D.N. Shcherbinin, A.S. Semikhin, Y. V Simakova, E.A. Tokarskaya, D.A. Egorova, M.M. Shmarov, N.A. Nikitenko, V.A. Gushchin, E.A. Smolyarchuk, S.K. Zyryanov, S. V Borisevich, B.S. Naroditsky, and A.L. Gintsburg. 2021. Safety and efficacy of an rAd26 and rAd5 vector-based heterologous prime-boost COVID-19 vaccine: an interim analysis of a randomised controlled phase 3 trial in Russia. Lancet. 397:671–681. doi:10.1016/s0140-6736(21)00234-8.

Lucas, C., P. Wong, J. Klein, T.B.R. Castro, J. Silva, M. Sundaram, M.K. Ellingson, T. Mao, J.E. Oh, B. Israelow, T. Takahashi, M. Tokuyama, P. Lu, A. Venkataraman, A. Park, S. Mohanty, H. Wang, A.L. Wyllie, C.B.F. Vogels, R. Earnest, S. Lapidus, I.M. Ott, A.J. Moore, M.C. Muenker, J.B. Fournier, M. Campbell, C.D. Odio, A. Casanovas-Massana, A. Obaid, A. Lu-Culligan, A. Nelson, A. Brito, A. Nunez, A. Martin, A. Watkins, B. Geng, C. Kalinich, C. Harden, C. Todeasa, C. Jensen, D. Kim, D. McDonald, D. Shepard, E. Courchaine, E.B. White, E. Song, E. Silva, E. Kudo, G. DeIuliis, H. Rahming, H.J. Park, I. Matos, J. Nouws, J. Valdez, J. Fauver, J. Lim, K.A. Rose, K. Anastasio, K. Brower, L. Glick, L. Sharma, L. Sewanan, L. Knaggs, M. Minasyan, M. Batsu, M. Petrone, M. Kuang, M. Nakahata, M. Campbell, M. Linehan, M.H. Askenase, M. Simonov, M. Smolgovsky, N. Sonnert, N. Naushad, P. Vijayakumar, R. Martinello, R. Datta, R. Handoko, S. Bermejo, S. Prophet, S. Bickerton, S. Velazquez, T. Alpert, T. Rice, W. Khoury-Hanold, X. Peng, Y. Yang, Y. Cao, Y. Strong, R. Herbst, A.C. Shaw, R. Medzhitov, W.L. Schulz, N.D. Grubaugh, C. Dela Cruz, S. Farhadian, A.I. Ko, et al. 2020. Longitudinal analyses reveal immunological misfiring in severe COVID-19. Nature. 584:463–469. doi:10.1038/s41586-020-2588-y.

Mahase, E. 2020. Covid-19: Moderna applies for US and EU approval as vaccine trial reports 94.1% efficacy. BMJ. 371. doi:10.1136/bmj.m4709.

Marshall, M. 2020. The lasting misery of coronavirus long-haulers. Nature. 585:339–341. doi:10.1038/d41586-020-02598-6.

Mateus, J., A. Grifoni, A. Tarke, J. Sidney, S.I. Ramirez, J.M. Dan, Z.C. Burger, S.A. Rawlings, D.M. Smith, E. Phillips, S. Mallal, M. Lammers, P. Rubiro, L. Quiambao, A. Sutherland, E.D. Yu, R. Da Silva Antunes, J. Greenbaum, A. Frazier, A.J. Markmann, L. Premkumar, A. De Silva, B. Peters, S. Crotty, A. Sette, and D. Weiskopf. 2020. Selective and cross-reactive SARS-CoV-2 T cell epitopes in unexposed humans. Science (80-.). 370:89–94. doi:10.1126/science.abd3871.

Mathew, D., J.R. Giles, A.E. Baxter, D.A. Oldridge, A.R. Greenplate, J.E. Wu, C. Alanio, L. Kuri-Cervantes, M.B. Pampena, K. D’Andrea, S. Manne, Z. Chen, Y.J. Huang, J.P. Reilly, A.R. Weisman, C.A.G. Ittner, O. Kuthuru, J. Dougherty, K. Nzingha, N. Han, J. Kim, A. Pattekar, E.C. Goodwin, E.M. Anderson, M.E. Weirick, S. Gouma, C.P. Arevalo, M.J. Bolton, F. Chen, S.F. Lacey, H. Ramage, S. Cherry, S.E. Hensley, S.A. Apostolidis, A.C. Huang, L.A. Vella, M.R. Betts, N.J. Meyer, E.J. Wherry, Z. Alam, M.M. Addison, K.T. Byrne, A. Chandra, H.C. Descamps, Y. Kaminskiy, J.T. Hamilton, J.H. Noll, D.K. Omran, E. Perkey, E.M. Prager, D. Pueschl, J.B. Shah, J.S. Shilan, and A.N. Vanderbeck. 2020. Deep immune profiling of COVID-19 patients reveals distinct immunotypes with therapeutic implications. Science (80-.). 369. doi:10.1126/SCIENCE.ABC8511.

McMahan, K., J. Yu, N.B. Mercado, C. Loos, L.H. Tostanoski, A. Chandrashekar, J. Liu, L. Peter, C. Atyeo, A. Zhu, E.A. Bondzie, G. Dagotto, M.S. Gebre, C. Jacob-Dolan, Z. Li, F. Nampanya, S. Patel, L. Pessaint, A. Van Ry, K. Blade, J. Yalley-Ogunro, M. Cabus, R. Brown, A. Cook, E. Teow, H. Andersen, M.G. Lewis, D.A. Lauffenburger, G. Alter, and D.H. Barouch. 2021. Correlates of protection against SARS-CoV-2 in rhesus macaques. Nature. 590:630–634. doi:10.1038/s41586-020-03041-6.

Meckiff, B.J., C. Ramírez-Suástegui, V. Fajardo, S.J. Chee, A. Kusnadi, H. Simon, S. Eschweiler, A. Grifoni, E. Pelosi, D. Weiskopf, A. Sette, F. Ay, G. Seumois, C.H. Ottensmeier, and P. Vijayanand. 2020. Imbalance of Regulatory and Cytotoxic SARS-CoV-2-Reactive CD4+ T Cells in COVID-19. Cell. 183:1340–1353.e16. doi:10.1016/j.cell.2020.10.001.

Nguyen, T.H.O., L.C. Rowntree, J. Petersen, B.Y. Chua, L. Hensen, L. Kedzierski, C.E. van de Sandt, P. Chaurasia, H.-X. Tan, J.R. Habel, W. Zhang, L. Allen, L. Earnest, K.Y. Mak, J.A. Juno, K. Wragg, F.L. Mordant, F. Amanat, F. Krammer, N.A. Mifsud, D.L. Doolan, K.L. Flanagan, S. Sonda, J. Kaur, L.M. Wakim, G.P. Westall, F. James, E. Mouhtouris, C.L. Gordon, N.E. Holmes, O.C. Smibert, J.A. Trubiano, A.C. Cheng, P. Harcourt, P. Clifton, J.C. Crawford, P.G. Thomas, A.K. Wheatley, S.J. Kent, J. Rossjohn, J. Torresi, and K. Kedzierska. 2021. CD8+ T cells specific for an immunodominant SARS-CoV-2 nucleocapsid epitope display high naïve precursor frequency and T cell receptor promiscuity. Immunity. 0. doi:10.1016/j.immuni.2021.04.009.

Noel, N., R. Peña, A. David, V. Avettand-Fenoel, I. Erkizia, E. Jimenez, C. Lecuroux, C. Rouzioux, F. Boufassa, G. Pancino, A. Venet, C. Van Lint, J. Martinez-Picado, O. Lambotte, A. Sáez-Cirión, and J.G. Prado. 2016. Long-Term Spontaneous Control of HIV-1 Is Related to Low Frequency of Infected Cells and Inefficient Viral Reactivation. J. Virol. 90:6148–6158. doi:10.1128/jvi.00419-16.

Oliver, S.E., J.W. Gargano, M. Marin, M. Wallace, K.G. Curran, M. Chamberland, N. McClung, D. Campos-Outcalt, R.L. Morgan, S. Mbaeyi, J.R. Romero, H.K. Talbot, G.M. Lee, B.P. Bell, and K. Dooling. 2020. The Advisory Committee on Immunization Practices’ Interim Recommendation for Use of Pfizer-BioNTech COVID-19 Vaccine — United States, December 2020. MMWR. Morb. Mortal. Wkly. Rep. 69:1922–1924. doi:10.15585/mmwr.mm6950e2.

Ou, X., Y. Liu, X. Lei, P. Li, D. Mi, L. Ren, L. Guo, R. Guo, T. Chen, J. Hu, Z. Xiang, Z. Mu, W. Chen, J. Chen, K. Hu, Q. Jin, J. Wang, and Z. Qian. 2020. Characterization of spike glycoprotein of SARS-CoV-2 on virus entry and its immune cross-reactivity with SARS-CoV. Nat. Commun. 11. doi:10.1038/s41467-020-15562-9.

Peng, Y., A.J. Mentzer, G. Liu, X. Yao, Z. Yin, D. Dong, W. Dejnirattisai, T. Rostron, P. Supasa, C. Liu, C. López-Camacho, J. Slon-Campos, Y. Zhao, D.I. Stuart, G.C. Paesen, J.M. Grimes, A.A. Antson, O.W. Bayfield, D.E.D.P. Hawkins, D.S. Ker, B. Wang, L. Turtle, K. Subramaniam, P. Thomson, P. Zhang, C. Dold, J. Ratcliff, P. Simmonds, T. de Silva, P. Sopp, D. Wellington, U. Rajapaksa, Y.L. Chen, M. Salio, G. Napolitani, W. Paes, P. Borrow, B.M. Kessler, J.W. Fry, N.F. Schwabe, M.G. Semple, J.K. Baillie, S.C. Moore, P.J.M. Openshaw, M.A. Ansari, S. Dunachie, E. Barnes, J. Frater, G. Kerr, P. Goulder, T. Lockett, R. Levin, Y. Zhang, R. Jing, L.P. Ho, T. Dong, P. Klenerman, A. McMichael, G. Ogg, J. Kenneth Baillie, R.J. Cornall, C.P. Conlon, G.R. Screaton, J. Mongkolsapaya, and J.C. Knight. 2020. Broad and strong memory CD4+ and CD8+ T cells induced by SARS-CoV-2 in UK convalescent individuals following COVID-19. Nat. Immunol. 21:1336–1345. doi:10.1038/s41590-020-0782-6.

Pereyra, F., X. Jia, P.J. McLaren, A. Telenti, P.I.W. de Bakker, B.D. Walker, S. Ripke, C.J. Brumme, S.L. Pulit, M. Carrington, C.M. Kadie, J.M. Carlson, D. Heckerman, R.R. Graham, R.M. Plenge, S.G. Deeks, L. Gianniny, G. Crawford, J. Sullivan, E. Gonzalez, L. Davies, A. Camargo, J.M. Moore, N. Beattie, S. Gupta, A. Crenshaw, N.P. Burtt, C. Guiducci, N. Gupta, X. Gao, Y. Qi, Y. Yuki, A. Piechocka-Trocha, E. Cutrell, R. Rosenberg, K.L. Moss, P. Lemay, J. O’leary, T. Schaefer, P. Verma, I. Toth, B. Block, B. Baker, A. Rothchild, J. Lian, J. Proudfoot, D.M.L. Alvino, S. Vine, M.M. Addo, T.M. Allen, M. Altfeld, M.R. Henn, S. Le Gall, H. Streeck, D.W. Haas, D.R. Kuritzkes, G.K. Robbins, R.W. Shafer, R.M. Gulick, C.M. Shikuma, R. Haubrich, S. Riddler, P.E. Sax, E.S. Daar, H.J. Ribaudo, B. Agan, S. Agarwal, R.L. Ahern, B.L. Allen, S. Altidor, E.L. Altschuler, S. Ambardar, K. Anastos, B. Anderson, V. Anderson, U. Andrady, D. Antoniskis, D. Bangsberg, D. Barbaro, W. Barrie, J. Bartczak, S. Barton, P. Basden, N. Basgoz, S. Bazner, N.C. Bellos, A.M. Benson, J. Berger, N.F. Bernard, A.M. Bernard, C. Birch, S.J. Bodner, R.K. Bolan, E.T. Boudreaux, M. Bradley, J.F. Braun, J.E. Brndjar, S.J. Brown, et al. 2010. The major genetic determinants of HIV-1 control affect HLA class I peptide presentation. Science (80-.). 330:1551–1557. doi:10.1126/science.1195271.

Pradenas, E., B. Trinité, V. Urrea, S. Marfil, C. Ávila-Nieto, M.L. Rodríguez de la Concepción, F. Tarrés-Freixas, S. Pérez-Yanes, C. Rovirosa, E. Ainsua-Enrich, J. Rodon, J. Vergara-Alert, J. Segalés, V. Guallar, A. Valencia, N. Izquierdo-Useros, R. Paredes, L. Mateu, A. Chamorro, M. Massanella, J. Carrillo, B. Clotet, and J. Blanco. 2021. Stable neutralizing antibody levels 6 months after mild and severe COVID-19 episodes. Med. 2:313–320.e4. doi:10.1016/j.medj.2021.01.005.

Rahman, M.M., and G. McFadden. 2006. Modulation of tumor necrosis factor by microbial pathogens. PLoS Pathog. 2:0066–0077. doi:10.1371/journal.ppat.0020004.

Reiss, S., A.E. Baxter, K.M. Cirelli, J.M. Dan, A. Morou, A. Daigneault, N. Brassard, G. Silvestri, J.P. Routy, C. Havenar-Daughton, S. Crotty, and D.E. Kaufmann. 2017. Comparative analysis of activation induced marker (AIM) assays for sensitive identification of antigen-specific CD4 T cells. PLoS One. 12. doi:10.1371/journal.pone.0186998.

Rha, M.S., H.W. Jeong, J.H. Ko, S.J. Choi, I.H. Seo, J.S. Lee, M. Sa, A.R. Kim, E.J. Joo, J.Y. Ahn, J.H. Kim, K.H. Song, E.S. Kim, D.H. Oh, M.Y. Ahn, H.K. Choi, J.H. Jeon, J.P. Choi, H. Bin Kim, Y.K. Kim, S.H. Park, W.S. Choi, J.Y. Choi, K.R. Peck, and E.C. Shin. 2021. PD-1-Expressing SARS-CoV-2-Specific CD8+ T Cells Are Not Exhausted, but Functional in Patients with COVID-19. Immunity. 54:44–52.e3. doi:10.1016/j.immuni.2020.12.002.

Roca, F.J., I. Mulero, A. López-Muñoz, M.P. Sepulcre, S.A. Renshaw, J. Meseguer, and V. Mulero. 2008. Evolution of the Inflammatory Response in Vertebrates: Fish TNF-α Is a Powerful Activator of Endothelial Cells but Hardly Activates Phagocytes. J. Immunol. 181:5071–5081. doi:10.4049/jimmunol.181.7.5071.

Ruiz-Riol, M., A. Llano, J. Ibarrondo, J. Zamarreño, K. Yusim, V. Bach, B. Mothe, S. Perez-Alvarez, M.A. Fernandez, G. Requena, M. Meulbroek, F. Pujol, A. Leon, P. Cobarsi, B.T. Korber, B. Clotet, C. Ganoza, J. Sanchez, J. Coll, and C. Brander. 2015. Alternative effector-function profiling identifies broad HIV-specific T-cell responses in highly HIV-exposed individuals who remain uninfected. J. Infect. Dis. 211:936–946. doi:10.1093/infdis/jiu534.

Rydyznski Moderbacher, C., S.I. Ramirez, J.M. Dan, A. Grifoni, K.M. Hastie, D. Weiskopf, S. Belanger, R.K. Abbott, C. Kim, J. Choi, Y. Kato, E.G. Crotty, C. Kim, S.A. Rawlings, J. Mateus, L.P.V. Tse, A. Frazier, R. Baric, B. Peters, J. Greenbaum, E. Ollmann Saphire, D.M. Smith, A. Sette, and S. Crotty. 2020. Antigen-Specific Adaptive Immunity to SARS-CoV-2 in Acute COVID-19 and Associations with Age and Disease Severity. Cell. 183:996–1012.e19. doi:10.1016/j.cell.2020.09.038.

Schulien, I., J. Kemming, V. Oberhardt, K. Wild, L.M. Seidel, S. Killmer, Sagar, F. Daul, M. Salvat Lago, A. Decker, H. Luxenburger, B. Binder, D. Bettinger, O. Sogukpinar, S. Rieg, M. Panning, D. Huzly, M. Schwemmle, G. Kochs, C.F. Waller, A. Nieters, D. Duerschmied, F. Emmerich, H.E. Mei, A.R. Schulz, S. Llewellyn-Lacey, D.A. Price, T. Boettler, B. Bengsch, R. Thimme, M. Hofmann, and C. Neumann-Haefelin. 2021. Characterization of pre-existing and induced SARS-CoV-2-specific CD8+ T cells. Nat. Med. 27:78–85. doi:10.1038/s41591-020-01143-2.

Schwarzkopf, S., A. Krawczyk, D. Knop, H. Klump, A. Heinold, F.M. Heinemann, L. Thümmler, C. Temme, M. Breyer, O. Witzke, U. Dittmer, V. Lenz, P.A. Horn, and M. Lindemann. 2021. Cellular Immunity in COVID-19 Convalescents with PCR-Confirmed Infection but with Undetectable SARS-CoV-2–Specific IgG. Emerg. Infect. Dis. 27:122–129. doi:10.3201/2701.203772.

Seder, R.A., P.A. Darrah, and M. Roederer. 2008. T-cell quality in memory and protection: Implications for vaccine design. Nat. Rev. Immunol. 8:247–258. doi:10.1038/nri2274.

Sekine, T., A. Perez-Potti, O. Rivera-Ballesteros, K. Strålin, J.B. Gorin, A. Olsson, S. Llewellyn-Lacey, H. Kamal, G. Bogdanovic, S. Muschiol, D.J. Wullimann, T. Kammann, J. Emgård, T. Parrot, E. Folkesson, M. Akber, L. Berglin, H. Bergsten, S. Brighenti, D. Brownlie, M. Butrym, B. Chambers, P. Chen, M.C. Jeannin, J. Grip, A.C. Gomez, L. Dillner, I.D. Lozano, M. Dzidic, M.F. Tullberg, A. Färnert, H. Glans, A. Haroun-Izquierdo, E. Henriksson, L. Hertwig, S. Kalsum, E. Kokkinou, E. Kvedaraite, M. Loreti, M. Lourda, K. Maleki, K.J. Malmberg, N. Marquardt, C. Maucourant, J. Michaelsson, J. Mjösberg, K. Moll, J. Muva, J. Mårtensson, P. Nauclér, A. Norrby-Teglund, L.P. Medina, B. Persson, L. Radler, E. Ringqvist, J.T. Sandberg, E. Sohlberg, T. Soini, M. Svensson, J. Tynell, R. Varnaite, A. Von Kries, C. Unge, O. Rooyackers, L.I. Eriksson, J.I. Henter, A. Sönnerborg, T. Allander, J. Albert, M. Nielsen, J. Klingström, S. Gredmark-Russ, N.K. Björkström, J.K. Sandberg, D.A. Price, H.G. Ljunggren, S. Aleman, and M. Buggert. 2020. Robust T Cell Immunity in Convalescent Individuals with Asymptomatic or Mild COVID-19. Cell. 183:158–168.e14. doi:10.1016/j.cell.2020.08.017.

Singh, A.K., S. Salwe, V. Padwal, S. Velhal, J. Sutar, S. Bhowmick, S. Mukherjee, V. Nagar, P. Patil, and V. Patel. 2020. Delineation of Homeostatic Immune Signatures Defining Viremic Non-progression in HIV-1 Infection. Front. Immunol. 11. doi:10.3389/fimmu.2020.00182.

Staines, H.M., D.E. Kirwan, D.J. Clark, E.R. Adams, Y. Augustin, R.L. Byrne, M. Cocozza, A.I. Cubas-Atienzar, L.E. Cuevas, M. Cusinato, B.M.O. Davies, M. Davis, P. Davis, A. Duvoix, N.M. Eckersley, D. Forton, A.J. Fraser, G. Garrod, L. Hadcocks, Q. Hu, M. Johnson, G.A. Kay, K. Klekotko, Z. Lewis, D.C. Macallan, J. Mensah-Kane, S. Menzies, I. Monahan, C.M. Moore, G. Nebe-Von-Caron, S.I. Owen, C. Sainter, A.A. Sall, J. Schouten, C.T. Williams, J. Wilkins, K. Woolston, J.R.A. Fitchett, S. Krishna, and T. Planche. 2021. Igg seroconversion and pathophysiology in severe acute respiratory syndrome coronavirus 2 infection. Emerg. Infect. Dis. 27:85–91. doi:10.3201/EID2701.203074.

Takahashi, T., M.K. Ellingson, P. Wong, B. Israelow, C. Lucas, J. Klein, J. Silva, T. Mao, J.E. Oh, M. Tokuyama, P. Lu, A. Venkataraman, A. Park, F. Liu, A. Meir, J. Sun, E.Y. Wang, A. Casanovas-Massana, A.L. Wyllie, C.B.F. Vogels, R. Earnest, S. Lapidus, I.M. Ott, A.J. Moore, K. Anastasio, M.H. Askenase, M. Batsu, H. Beatty, S. Bermejo, S. Bickerton, K. Brower, M.L. Bucklin, S. Cahill, M. Campbell, Y. Cao, E. Courchaine, R. Datta, G. DeIuliis, B. Geng, L. Glick, R. Handoko, C. Kalinich, W. Khoury-Hanold, D. Kim, L. Knaggs, M. Kuang, E. Kudo, J. Lim, M. Linehan, A. Lu-Culligan, A.A. Malik, A. Martin, I. Matos, D. McDonald, M. Minasyan, S. Mohanty, M.C. Muenker, N. Naushad, A. Nelson, J. Nouws, M. Nunez-Smith, A. Obaid, I. Ott, H.J. Park, X. Peng, M. Petrone, S. Prophet, H. Rahming, T. Rice, K.A. Rose, L. Sewanan, L. Sharma, D. Shepard, E. Silva, M. Simonov, M. Smolgovsky, E. Song, N. Sonnert, Y. Strong, C. Todeasa, J. Valdez, S. Velazquez, P. Vijayakumar, H. Wang, A. Watkins, E.B. White, Y. Yang, A. Shaw, J.B. Fournier, C.D. Odio, S. Farhadian, C. Dela Cruz, N.D. Grubaugh, W.L. Schulz, A.M. Ring, A.I. Ko, S.B. Omer, and A. Iwasaki. 2020. Sex differences in immune responses that underlie COVID-19 disease outcomes. Nature. 588:315–320. doi:10.1038/s41586-020-2700-3.

Tegally, H., E. Wilkinson, M. Giovanetti, A. Iranzadeh, V. Fonseca, J. Giandhari, D. Doolabh, S. Pillay, E.J. San, N. Msomi, K. Mlisana, A. von Gottberg, S. Walaza, M. Allam, A. Ismail, T. Mohale, A.J. Glass, S. Engelbrecht, G. Van Zyl, W. Preiser, F. Petruccione, A. Sigal, D. Hardie, G. Marais, M. Hsiao, S. Korsman, M.-A. Davies, L. Tyers, I. Mudau, D. York, C. Maslo, D. Goedhals, S. Abrahams, O. Laguda-Akingba, A. Alisoltani-Dehkordi, A. Godzik, C.K. Wibmer, B.T. Sewell, J. Lourenço, L.C.J. Alcantara, S.L. Kosakovsky Pond, S. Weaver, D. Martin, R.J. Lessells, J.N. Bhiman, C. Williamson, and T. de Oliveira. 2021. Emergence of a SARS-CoV-2 variant of concern with mutations in spike glycoprotein. Nature. doi:10.1038/s41586-021-03402-9.

Thieme, C.J., M. Anft, K. Paniskaki, A. Blazquez-Navarro, A. Doevelaar, F.S. Seibert, B. Hoelzer, M.J. Konik, M.M. Berger, T. Brenner, C. Tempfer, C. Watzl, T.L. Meister, S. Pfaender, E. Steinmann, S. Dolff, U. Dittmer, T.H. Westhoff, O. Witzke, U. Stervbo, T. Roch, and N. Babel. 2020. Robust T Cell Response Toward Spike, Membrane, and Nucleocapsid SARS-CoV-2 Proteins Is Not Associated with Recovery in Critical COVID-19 Patients. Cell Reports Med. 1. doi:10.1016/j.xcrm.2020.100092.

Trinité, B., F. Tarrés-Freixas, J. Rodon, E. Pradenas, V. Urrea, S. Marfil, M.L. Rodríguez de la Concepción, C. Ávila-Nieto, C. Aguilar-Gurrieri, A. Barajas, R. Ortiz, R. Paredes, L. Mateu, A. Valencia, V. Guallar, L. Ruiz, E. Grau, M. Massanella, J. Puig, A. Chamorro, N. Izquierdo-Useros, J. Segalés, B. Clotet, J. Carrillo, J. Vergara-Alert, and J. Blanco. 2021. SARS-CoV-2 infection elicits a rapid neutralizing antibody response that correlates with disease severity. Sci. Rep. 11. doi:10.1038/s41598-021-81862-9.

Vahidy, F.S., A.P. Pan, H. Ahnstedt, Y. Munshi, H.A. Choi, Y. Tiruneh, K. Nasir, B.A. Kash, J.D. Andrieni, and L.D. McCullough. 2021. Sex differences in susceptibility, severity, and outcomes of coronavirus disease 2019: Cross-sectional analysis from a diverse US metropolitan area. PLoS One. 16:e0245556. doi:10.1371/journal.pone.0245556.

Voysey, M., S.A. Costa Clemens, S.A. Madhi, L.Y. Weckx, P.M. Folegatti, P.K. Aley, B. Angus, V.L. Baillie, S.L. Barnabas, Q.E. Bhorat, S. Bibi, C. Briner, P. Cicconi, E.A. Clutterbuck, A.M. Collins, C.L. Cutland, T.C. Darton, K. Dheda, C. Dold, C.J.A. Duncan, K.R.W. Emary, K.J. Ewer, A. Flaxman, L. Fairlie, S.N. Faust, S. Feng, D.M. Ferreira, A. Finn, E. Galiza, A.L. Goodman, C.M. Green, C.A. Green, M. Greenland, C. Hill, H.C. Hill, I. Hirsch, A. Izu, D. Jenkin, C.C.D. Joe, S. Kerridge, A. Koen, G. Kwatra, R. Lazarus, V. Libri, P.J. Lillie, N.G. Marchevsky, R.P. Marshall, A.V.A. Mendes, E.P. Milan, A.M. Minassian, A. McGregor, Y.F. Mujadidi, A. Nana, S.D. Padayachee, D.J. Phillips, A. Pittella, E. Plested, K.M. Pollock, M.N. Ramasamy, A.J. Ritchie, H. Robinson, A. V Schwarzbold, A. Smith, R. Song, M.D. Snape, E. Sprinz, R.K. Sutherland, E.C. Thomson, M.E. Török, M. Toshner, D.P.J. Turner, J. Vekemans, T.L. Villafana, T. White, C.J. Williams, A.D. Douglas, A.V.S. Hill, T. Lambe, S.C. Gilbert, A.J. Pollard, and Oxford COVID Vaccine Trial Group. 2021. Single-dose administration and the influence of the timing of the booster dose on immunogenicity and efficacy of ChAdOx1 nCoV-19 (AZD1222) vaccine: a pooled analysis of four randomised trials. Lancet (London, England). 397:881–891. doi:10.1016/S0140-6736(21)00432-3.

Wilk, A.J., A. Rustagi, N.Q. Zhao, J. Roque, G.J. Martínez-Colón, J.L. McKechnie, G.T. Ivison, T. Ranganath, R. Vergara, T. Hollis, L.J. Simpson, P. Grant, A. Subramanian, A.J. Rogers, and C.A. Blish. 2020. A single-cell atlas of the peripheral immune response in patients with severe COVID-19. Nat. Med. 26:1070–1076. doi:10.1038/s41591-020-0944-y.

Zhao, Q., M. Meng, R. Kumar, Y. Wu, J. Huang, Y. Deng, Z. Weng, and L. Yang. 2020. Lymphopenia is associated with severe coronavirus disease 2019 (COVID-19) infections: A systemic review and meta-analysis. Int. J. Infect. Dis. 96:131–135. doi:10.1016/j.ijid.2020.04.086.

Zheng, H.-Y., M. Zhang, C.-X. Yang, N. Zhang, X.-C. Wang, X.-P. Yang, X.-Q. Dong, and Y.-T. Zheng. 2020. Elevated exhaustion levels and reduced functional diversity of T cells in peripheral blood may predict severe progression in COVID-19 patients. Cell. Mol. Immunol. 17:541–543. doi:10.1038/s41423-020-0401-3.

